# Leukemia in Sprague-Dawley Rats Exposed Long-term from Prenatal Life to Glyphosate and Glyphosate-Based Herbicides

**DOI:** 10.1101/2023.11.14.566013

**Authors:** Simona Panzacchi, Eva Tibaldi, Luana De Angelis, Laura Falcioni, Federica Gnudi, Martina Iuliani, Marco Manservigi, Fabiana Manservisi, Isabella Manzoli, Ilaria Menghetti, Rita Montella, Roberta Noferini, Daria Sgargi, Valentina Strollo, Michael Antoniou, Jia Chen, Giovanni Dinelli, Stefano Lorenzetti, Robin Mesnage, Andrea Vornoli, Melissa J. Perry, Philip J. Landrigan, Fiorella Belpoggi, Daniele Mandrioli

## Abstract

**Background:** Glyphosate-based herbicides (GBHs) are the world’s most widely used weed control agents. There has been intense and increasing public health concern about glyphosate and GBHs since the International Agency for Research on Cancer classified glyphosate as a probable human carcinogen in 2015.

**Aims:** To further study the health effects of glyphosate and GBHs, the Ramazzini Institute, in collaboration with an international network of institutes and universities, has launched the Global Glyphosate Study (GGS), the most comprehensive toxicological study ever performed on these compounds. The GGS is an integrated study designed to test a wide range of toxicological outcomes including carcinogenicity, neurotoxicity, multi-generational effects, organ toxicity, endocrine disruption and prenatal developmental toxicity. The present study reports the first definitive results on leukemia incidence and mortality from the carcinogenicity arm of the GGS.

**Method:** Glyphosate and two GBHs, Roundup Bioflow (MON 52276) used in the European Union (EU) and RangerPro (EPA 524-517) used in the U.S., were administered long-term to Sprague-Dawley (SD) rats beginning in prenatal life until 104 weeks of age via drinking water at doses of 0.5, 5, and 50 mg/kg body weight/day. This dose range encompasses both the EU Acceptable Daily Intake (ADI) and the EU No Observed Adverse Effect Level (NOAEL) for glyphosate. Each experimental group was composed of 51 males and 51 females, the total number animals were 1020 (510 males and 510 females).

**Results:** In the animals exposed to glyphosate, a significantly increased trend in incidence of lymphoblastic leukemia was observed in males. In the Roundup Bioflow-treated animals, significantly increased trends were observed in incidence of lymphoblastic leukemia (males and females), monocytic leukemia (males), total myeloid leukemia (males), and all leukemias combined (males and females). In the RangerPro-treated animals, significantly increased trends were observed in incidence of lymphoblastic leukemia (males and females), monocytic leukemia (males) and all leukemias combined (males). 43% of leukemias deaths in the glyphosate and GBHs treated groups occurred before the first year of age (52 weeks).

**Conclusions:** Glyphosate and GBHs at exposure levels corresponding to the EU ADI and the EU NOAEL caused significant, dose-related increased trends in incidence of leukemia, a very rare malignancy, in SD rats. Notably, about half of the leukemia deaths seen in the glyphosate and GBH groups occurred at less than one year of age, comparable to less than 35-40 years of age in humans.

## Introduction

Glyphosate [IUPAC chemical name N-(phosphonomethyl)glycine], is a broad-spectrum herbicide used for non-selective weed control both in conventional agriculture and non-agricultural settings. It acts by inhibiting the enzyme 5-enolpyruvylshikimate-3-phosphate synthase, which is involved in the synthesis of essential amino acids in plants. Glyphosate’s extensive use reflects its effectiveness in controlling a wide range of both broadleaf and grassy weeds. Glyphosate and its metabolites persist in food (Rodrigues and de Souza 2018; Wang et al. 1994), water (Alferness and Iwata 1994), and dust (Navarro et al. 2023), indicating potential for most populations exposure (Silva et al. 2023).

In March 2015, the International Agency for Research on Cancer (IARC), the cancer agency of the World Health Organization (WHO), classified glyphosate as *probably carcinogenic to humans* (Group 2A) based on *limited evidence of carcinogenicity* in humans, based on non-Hodgkin lymphoma, and *sufficient evidence of carcinogenicity in rodents* (IARC 2015). The IARC working group also concluded, based on publicly available literature, that there was strong evidence including in human cells in vitro that glyphosate is genotoxic and that it induced oxidative stress, which are two key characteristics of carcinogens (Smith et al. 2016). Exposure to GBHs may be more toxic than exposure to glyphosate alone (Manservisi et al. 2019; Mao et al. 2018; R. Mesnage et al. 2022b; Truzzi et al. 2021). GBHs contain not only glyphosate, but also co-formulants with no direct herbicidal action that facilitate glyphosate penetration into tissues through the waxy surface of plant leaves (Mesnage et al. 2019). In 2016 the European Union (EU) banned polyoxyethylene tallow amine (POEA)-type co-formulants from use in GBHs, but these formulations are still in use outside the EU (Langrand et al. 2020; Mesnage and Antoniou 2018; Mesnage et al. 2019). Following the EU phase-out of POEA, GBH manufacturers have turned to other surfactants.

Recent epidemiologic studies have expanded knowledge of the human carcinogenicity of glyphosate and GBHs. A recent meta-analysis reports that GBHs are associated with a 41% increased relative risk of non-Hodgkin lymphoma among highly exposed individuals (Zhang et al. 2019). Skidmore et al. showed a significant temporal and geographical relationship between expansion of soy cultivation in the Brazilian Amazon and Cerrado regions and deaths from acute lymphoblastic leukemia in children, one of the most common pediatric blood-borne cancer (Skidmore et al. 2023). Hardell et al. published the results of a pooled analysis of three Swedish case-control studies including 1425 cases and 2157 controls, examining exposures to phenoxyacetic acids and glyphosate in relation to non-Hodgkin lymphoma (Hardell et al. 2023). This analysis confirmed an association between non-Hodgkin lymphoma and exposure to these herbicides (Hardell et al. 2023). Another study, focused on fenceline exposure to agricultural pesticides and risk of childhood leukemia in an Italian community, reported an increasing risk of leukemia in children residing close to arable crops exposed to a mixture of glyphosate and other pesticides (Malagoli et al. 2016). In the updated follow-up report of the United States Agricultural Health Study, a large prospective cohort investigating cancer incidence through 2012 (North Carolina)/2013 (Iowa) with 7290 incident cancer cases, evidence of increased risk of acute myeloid leukemia was observed among the group with highest exposure to glyphosate (Andreotti et al. 2018).

Animal carcinogenicity studies support this expanding epidemiologic literature by showing glyphosate’s ability to cause multiple types of cancer, especially malignant lymphoma (Portier 2020).

In 2019, the Ramazzini Institute launched the Global Glyphosate Study (GGS), an independent, collaborative, multi-institutional study with the aim of providing the most comprehensive toxicological evaluation of glyphosate and GBHs (glyphosatestudy.org). The GGS is an integrated study designed to test a wide range of toxicological outcomes including carcinogenicity, neurotoxicity, multi-generational effects, organ toxicity, endocrine disruption and prenatal developmental toxicity.

This report presents the first definitive results on leukemia incidence and mortality from the carcinogenicity arm of the GGS, in which glyphosate and two commercial formulations, namely the EU representative formulation Roundup Bioflow (MON 52276) and the US formulation RangerPro (EPA 524-517), were administered long-term to SD rats beginning prenatal life via drinking water at 0.5, 5, and 50 mg/kg bw/day. This dose range encompasses both the EU acceptable daily intake (ADI) and the EU no-observed-adverse-effect level (NOAEL) for glyphosate. RangerPro contains POEA surfactants (Robin Mesnage et al. 2022), while Roundup Bioflow does not contain POEA (Mesnage et al. 2021b), however the complete co-formulant profile remains unknown. Evaluations of other neoplastic lesions from the carcinogenicity arm of the GGS are currently ongoing.

## Materials and Methods

### Chemicals

The test substances were administered to female and male SD rats in drinking water:

- Active ingredient Glyphosate CAS 38641-94-0 [N-(phosphonomethyl)glycine, purity 96%, Sigma-Aldrich, Milan, Italy].
- The commercial formulation Roundup Bioflow [MON 52276, containing 360 g/L of glyphosate acid in the form of 486 g/L isopropylamine salts of glyphosate (41.5%)].
- The commercial formulation RangerPro [EPA registration n. 524-517, containing 356 g/L of glyphosate acid in the form of 480 g/L isopropylamine salts of glyphosate (41%)].

Solutions of these three test substances were prepared by the addition of appropriate volume of tap drinking water, to obtain the corresponding concentration of glyphosate.

### Animals and experimental design

The entire animal experiment was performed in accord with the Italian laws regulating the use and treatment of animals for scientific purposes (Legislative Decree No. 26, 2014. Implementation of the directive n. 2010/63/EU on the protection of animals used for scientific purposes. G.U. General Series, n. 61, March 14th, 2014). All animal study procedures were performed at the Cesare Maltoni Cancer Research Centre/Ramazzini Institute (CMCRC/RI), Bentivoglio, Italy. The study protocol was approved by the Ethical Committee of the Ramazzini Institute. The protocol of the experiment was also approved and formally authorized by the ad hoc commission of the Italian Ministry of Health (ministerial approval n. 945/2018-PR).

The CMCRC/RI animal breeding facility was the supplier for the SD rats. Female breeder SD rats were placed individually in polycarbonate cages (42x26x18cm; Tecniplast Buguggiate, Varese, Italy) with a single unrelated male until evidence of copulation was observed. After mating, the male was removed and females were housed separately during gestation and delivery. Newborns were housed with their dams until weaning. Weaned offspring were co-housed, by sex and treatment group, no more than 3 each per cage. Cages were identified by a card indicating: study protocol code, experimental and pedigree numbers, and dosage group. A shallow layer of white fir wood shavings served as bedding (supplier: Giuseppe Bordignon, Treviso, Italy).

Analyses of chemical characteristics (pH, ashes, dry weight, and specific weight) and possible contamination (metals, aflatoxin, polychlorinated biphenyls, organophosphorus and organochlorine pesticides) of the bedding was performed by CONSULAB srl Laboratories (Treviso, Italy). Both feed and water were analyzed to identify possible chemical or microbiological contaminants or impurities.

The cages were placed on racks at room temperature of 22°C ± 3°C and 50% ± 20% of relative humidity. Daily checks on temperature and humidity were performed. The light was artificial, and a light/dark cycle of 12 hours was maintained.

Sprague-Dawley rat dams (F0) and relative pups (F1) were treated with either glyphosate or Roundup Bioflow or RangerPro diluted in drinking water to achieve the desired glyphosate concentration. Three doses were selected for each test substance, 5, 50 and 500 mg/l of glyphosate in drinking water, based on mean water consumption and a mean body weight, corresponding to 0.5, 5 and 50 mg/kg bw/day. Control animals were administered drinking water *ad libitum*. The total number of experimental groups were 10: control group (drinking water), glyphosate groups (3 doses), Roundup Bioflow groups (3 doses), RangerPro groups (3 doses). F0 animals were randomly distributed in order to have 1 male and 1 female per litter per group and mated outbred. The F0 female breeders received the treatment through drinking water from gestation day (GD) 6 to the end of lactation, while the offspring (F1) continued to be exposed after weaning until study termination at 104 weeks of age. After weaning, F1 animals were randomly distributed in order to have no more than 2 males and 2 females per litter per group. Each F1 experimental group was composed of 51 males and 51 females, belonging to the same treatment group of their exposed dams. The total numbers of animals was 1020 (510 males and 510 females) (Table 1-3).

**Table 1.** Incidence of leukemia in carcinogenicity bioassays on glyphosate administered in drinking water *ad libitum* at three doses and in untreated controls to male (M) and female (F) Sprague-Dawley rats.

| Group No. | Dose (mg/kg bw/day of Glyphosate) | Animals |  | Lymphoblastic leukaemias |  |  | Monocytic leukaemias |  | Myeloid leukemias |  | Total lymphoid leukaemias |  |  | Total myeloid leukaemias |  | All leukaemias |  |
| --- | --- | --- | --- | --- | --- | --- | --- | --- | --- | --- | --- | --- | --- | --- | --- | --- | --- |
|  |  | Sex | No. | No. | % | P-value | No. | % | No. | % | No. | % | P-value | No. | % | No. | % |
| I | 0 (control) | M | 51 | 0 | 0.0 | #0.0419<br>*0.0335 | 0 | 0.0 | 0 | 0.0 | 0 | 0.0 | #0.0419<br>*0.0335 | 0 | 0.0 | 0 | 0.0 |
|  |  | F | 51 | 0 | 0.0 |  | 0 | 0.0 | 0 | 0.0 | 0 | 0.0 |  | 0 | 0.0 | 0 | 0.0 |
|  |  | M+F | 102 | 0 | 0.0 | #0.0421<br>*0.0372 | 0 | 0.0 | 0 | 0.0 | 0 | 0.0 | #0.0421<br>*0.0372 | 0 | 0.0 | 0 | 0.0 |
| II | 0.5 Glyphosate | M | 51 | 0 | 0.0 |  | 1 | 2.0 | 0 | 0.0 | 0 | 0.0 |  | 1 | 2.0 | 1 | 2.0 |
|  |  | F | 51 | 0 | 0.0 |  | 0 | 0.0 | 1 | 2.0 | 0 | 0.0 |  | 1 | 2.0 | 1 | 2.0 |
|  |  | M+F | 102 | 0 | 0.0 |  | 1 | 1.0 | 1 | 1.0 | 0 | 0.0 |  | 2 | 2.0 | 2 | 2.0 |
| III | 5 Glyphosate | M | 51 | 0 | 0.0 |  | 1 | 2.0 | 0 | 0.0 | 0 | 0.0 |  | 1 | 2.0 | 1 | 2.0 |
|  |  | F | 51 | 0 | 0.0 |  | 0 | 0.0 | 0 | 0.0 | 0 | 0.0 |  | 0 | 0.0 | 0 | 0.0 |
|  |  | M+F | 102 | 0 | 0.0 |  | 1 | 1.0 | 0 | 0.0 | 0 | 0.0 |  | 1 | 1.0 | 1 | 1.0 |
| IV | 50 Glyphosate | M | 51 | 1 | 2.0 |  | 0 | 0.0 | 0 | 0.0 | 1 | 2.0 |  | 0 | 0.0 | 1 | 2.0 |
|  |  | F | 51 | 0 | 0.0 |  | 0 | 0.0 | 0 | 0.0 | 0 | 0.0 |  | 0 | 0.0 | 0 | 0.0 |
|  |  | M+F | 102 | 1 | 1.0 |  | 0 | 0.0 | 0 | 0.0 | 1 | 1.0 |  | 0 | 0.0 | 1 | 1.0 |

**Table 2.** Incidence of leukemia in carcinogenicity bioassays on Roundup Bioflow administered in drinking water ad libitum at three doses and in untreated controls to male (M) and female (F) Sprague-Dawley rats.

| Group No. | Dose (mg/kg bw/day of Glyphosate) | Animals |  | Lymphoblastic leukaemias |  |  | Monocytic leukaemias |  |  | Myeloid leukemias |  | Total lymphoid leukaemias |  |  | Total myeloid leukaemias |  |  | All leukaemias |  |  |
| --- | --- | --- | --- | --- | --- | --- | --- | --- | --- | --- | --- | --- | --- | --- | --- | --- | --- | --- | --- | --- |
|  |  | Sex | No. | No. | % | P-value | No. | % | P-value | No. | % | No. | % | P-value | No. | % | P-value | No. | % | P-value |
| I | 0 (control) | M | 51 | 0 | 0.0 | #0.0419<br>*0.0425 | 0 | 0.0 | #0.0419<br>*0.0425 | 0 | 0.0 | 0 | 0.0 | #0.0419<br>*0.0425 | 0 | 0.0 | #0.0419<br>*0.0425 | 0 | 0.0 | #0.0071<br>*0.0083 |
|  |  | F | 51 | 0 | 0.0 | #0.0419<br>*0.0439 | 0 | 0.0 |  | 0 | 0.0 | 0 | 0.0 | #0.0419<br>*0.0439 | 0 | 0.0 |  | 0 | 0.0 | #0.0419<br>*0.0439 |
|  |  | M+F | 102 | 0 | 0.0 | #0.0072<br>*0.0078 | 0 | 0.0 | #0.0421<br>*0.0432 | 0 | 0.0 | 0 | 0.0 | #0.0072<br>*0.0078 | 0 | 0.0 | #0.0421<br>*0.0432 | 0 | 0.0 | #0.0014<br>*0.0016 |
| V | 0.5 Roundup Bioflow | M | 51 | 0 | 0.0 |  | 0 | 0.0 |  | 0 | 0.0 | 0 | 0.0 |  | 0 | 0.0 |  | 0 | 0.0 |  |
|  |  | F | 51 | 0 | 0.0 |  | 0 | 0.0 |  | 0 | 0.0 | 0 | 0.0 |  | 0 | 0.0 |  | 0 | 0.0 |  |
|  |  | M+F | 102 | 0 | 0.0 |  | 0 | 0.0 |  | 0 | 0.0 | 0 | 0.0 |  | 0 | 0.0 |  | 0 | 0.0 |  |
| VI | 5 Roundup Bioflow | M | 51 | 0 | 0.0 |  | 0 | 0.0 |  | 0 | 0.0 | 0 | 0.0 |  | 0 | 0.0 |  | 0 | 0.0 |  |
|  |  | F | 51 | 0 | 0.0 |  | 0 | 0.0 |  | 0 | 0.0 | 0 | 0.0 |  | 0 | 0.0 |  | 0 | 0.0 |  |
|  |  | M+F | 102 | 0 | 0.0 |  | 0 | 0.0 |  | 0 | 0.0 | 0 | 0.0 |  | 0 | 0.0 |  | 0 | 0.0 |  |
| VII | 50 Roundup Bioflow | M | 51 | 1 | 2.0 |  | 1 | 2.0 |  | 0 | 0.0 | 1 | 2.0 |  | 1 | 2.0 |  | 2 | 3.9 |  |
|  |  | F | 51 | 1 | 2.0 |  | 0 | 0.0 |  | 0 | 0.0 | 1 | 2.0 |  | 0 | 0.0 |  | 1 | 2.0 |  |
|  |  | M+F | 102 | 2 | 2.0 |  | 1 | 1.0 |  | 0 | 0.0 | 2 | 2.0 |  | 1 | 1.0 |  | 3 | 2.9 |  |

**Table 3.** Incidence of leukemia in carcinogenicity bioassays on RangerPro administered in drinking water *ad libitum* at three doses and in untreated controls to male (M) and female (F) Sprague-Dawley rats.

| Group No. | Dose (mg/kg bw/day of Glyphosate) | Animals |  | Lymphoblastic leukaemias |  |  | Monocytic leukaemias |  |  | Myeloid leukemias |  | Total lymphoid leukaemias |  |  | Total myeloid leukaemias |  | All leukaemias |  |  |
| --- | --- | --- | --- | --- | --- | --- | --- | --- | --- | --- | --- | --- | --- | --- | --- | --- | --- | --- | --- |
|  |  | Sex | No. | No. | % | P-value | No. | % | P-value | No. | % | No. | % | P-value | No. | % | No. | % | P-value |
| I | 0 (control) | M | 51 | 0 | 0.0 | #0.0071<br>*0.0092 | 0 | 0.0 | #0.0419<br>*0.0459 | 0 | 0.0 | 0 | 0.0 | #0.0071<br>*0.0092 | 0 | 0.0 | 0 | 0.0 | #0.0313<br>*0.0371 |
|  |  | F | 51 | 0 | 0.0 | #0.0419<br>*0.0411 | 0 | 0.0 |  | 0 | 0.0 | 0 | 0.0 | #0.0419<br>*0.0411 | 0 | 0.0 | 0 | 0.0 |  |
|  |  | M+F | 102 | 0 | 0.0 | #0.0014<br>*0.0016 | 0 | 0.0 |  | 0 | 0.0 | 0 | 0.0 | #0.0014<br>*0.0016 | 0 | 0.0 | 0 | 0.0 | #0.0195<br>*0.0218 |
| VIII | 0.5 RangerPro | M | 51 | 0 | 0.0 |  | 0 | 0.0 |  | 1 | 2.0 | 0 | 0.0 |  | 1 | 2.0 | 1 | 2.0 |  |
|  |  | F | 51 | 0 | 0.0 |  | 0 | 0.0 |  | 0 | 0.0 | 0 | 0.0 |  | 0 | 0.0 | 0 | 0.0 |  |
|  |  | M+F | 102 | 0 | 0.0 |  | 0 | 0.0 |  | 1 | 1.0 | 0 | 0.0 |  | 1 | 1.0 | 1 | 1.0 |  |
| IX | 5 RangerPro | M | 51 | 0 | 0.0 |  | 0 | 0.0 |  | 1 | 2.0 | 0 | 0.0 |  | 1 | 2.0 | 1 | 2.0 |  |
|  |  | F | 51 | 0 | 0.0 |  | 1 | 2.0 |  | 0 | 0.0 | 0 | 0.0 |  | 1 | 2.0 | 1 | 2.0 |  |
|  |  | M+F | 102 | 0 | 0.0 |  | 1 | 1.0 |  | 1 | 1.0 | 0 | 0.0 |  | 2 | 2.0 | 2 | 2.0 |  |
| X | 50 RangerPro | M | 51 | 2 | 3.9 |  | 1 | 2.0 |  | 0 | 0.0 | 2 | 3.9 |  | 1 | 2.0 | 3 | 5.9 |  |
|  |  | F | 51 | 1 | 2.0 |  | 0 | 0.0 |  | 0 | 0.0 | 1 | 2.0 |  | 0 | 0.0 | 1 | 2.0 |  |
|  |  | M+F | 102 | 3 | 2.9 |  | 1 | 1.0 |  | 0 | 0.0 | 3 | 2.9 |  | 1 | 1.0 | 4 | 3.9 |  |

### Necropsy and histopathology

The *in vivo* phase ended at 104 weeks of age. Upon death, all animals underwent a complete necropsy. Gross necropsy, performed on all animals, included an initial physical examination of the external surfaces and all orifices followed by an internal examination of tissues and organs *in situ*. Histopathology was performed on the following organs and tissues of each animal from each group: skin and subcutaneous tissue, mammary gland (4 sites: axillary and inguinal, right and left), brain with cerebellum and medulla/pons, pituitary gland, salivary glands, Harderian glands, tongue, esophagus, thyroid and parathyroid, thymus and mediastinal lymph nodes, trachea, lungs, heart, liver (2 lobes for histopathology), spleen, pancreas, kidneys, adrenal glands, stomach (forestomach and glandular stomach), small intestine, large intestine (with the Peyer’s patches), bladder, uterus (including cervix), ovaries, vagina, testes and epididymis, seminal vesicles and coagulating glands, prostate, subcutaneous lymph nodes, mesenteric lymph nodes, all gross lesions and other tissues only if anomalies were present. After fixation, samples were trimmed, processed, embedded in paraffin wax, sectioned to a thickness of 4-5 μm and then processed in alcohol-xylene series and stained with hematoxylin and eosin for microscopic evaluation. Histopathology evaluation was performed by at least two pathologists and all lesions were peer-reviewed. The pathological evaluations were performed according to the procedures of the International Harmonization of Nomenclature and Diagnostic Criteria (INHAND)(Willard-Mack et al. 2019).

### Statistical methods

For all analyses, p values ≤0.05 were considered statistically significant. Data were collected from both sexes and analyses were conducted separately.

### Survival analysis

The probability of survival was estimated by the product-limit procedure of Kaplan and Meier and was presented graphically (Kaplan and Meier 1958). All animals were considered in the analysis, including those dying from unnatural causes within the observation period (humanitarian sacrifices) and those surviving to the end of the observation period (terminated via final sacrifice; their survival time was fixed at 104 weeks).

Dose-related differences and trends were assessed using a log-rank test, first on all experimental groups together, and then separately by exposure group (Glyphosate vs Control, Roundup Bioflow vs Control, RangerPro vs Control). For the subgroup of animals bearing any type of leukemia, dose-related trends were examined with Tarone and Ware’s life-table test (Tarone 1975). All reported p values for the survival analyses are two-sided.

### Analysis of Neoplastic Lesion Incidence

Incidence of leukemias was calculated as the number of animals with leukemic lesions divided by the total number of animals examined microscopically. Tests of significance included pairwise comparisons of each exposed group with the control group using a one-tailed Fisher’s exact test (one-sided results were considered, because it is well established that only an increase in the disease incidences can be expected from the exposure, and incidences of leukemia in the control group are almost always 0). Furthermore, the Cochran-Armitage trend test was performed to test for dose-related trends within different exposures (glyphosate vs Control, Roundup Bioflow vs Control, RangerPro vs Control). Reported p values are one-sided.

Survival differences among groups were considered in assessing incidence of neoplastic lesions. In this assessment, the Poly-k test was used (Bailer and Portier 1988; Piegorsch and Bailer 1997; Portier and Bailer 1989; Portier et al. 1986). This test is a survival-adjusted quantal-response procedure that modifies the Cochran-Armitage linear trend test to account for survival differences by assigning a reduced risk of neoplasm, proportional to a power of the fraction of time on study, to only those lesion-free animals that did not reach terminal euthanasia. More specifically, this method modifies the denominator in the quantal estimate of lesion incidence to approximate more closely the total number of animal years at risk. Each animal is assigned a risk weight: this value is 1 if the animal had a lesion at the site under consideration or if it survived until terminal euthanasia; if the animal died before terminal sacrifice and did not have a lesion at the site under consideration, its risk weight is the fraction of the entire study time that it survived, raised to the k-th power. A value of k = 3 was used in the analysis of leukemias, as recommended by Bailer and Portier (Bailer and Portier 1988; Piegorsch and Bailer 1997). Reported p values for these analyses are one-sided.

Statistical analyses were performed using the German Cancer Research Center software (https://biostatistics.dkfz.de/) and Stata 18 software (StataCorp 2023. Stata Statistical Software: Release 18. College Station, TX: StataCorp LLC).

## Results

### In vivo end points

During gestation and lactation, the *in vivo* phase was carried out without showing toxic and/or adverse effects. Litter sizes were similar among groups. Mean body weights of dams and pups was in the range of 10% variability relative to controls; food consumption was in the range of 20% variability relative to controls; water consumption was in the range of 20% variability relative to control. After weaning, body weights, water and feed consumption were homogenous among groups in both females and males. (**Figure 1**).

**Figure 1.**
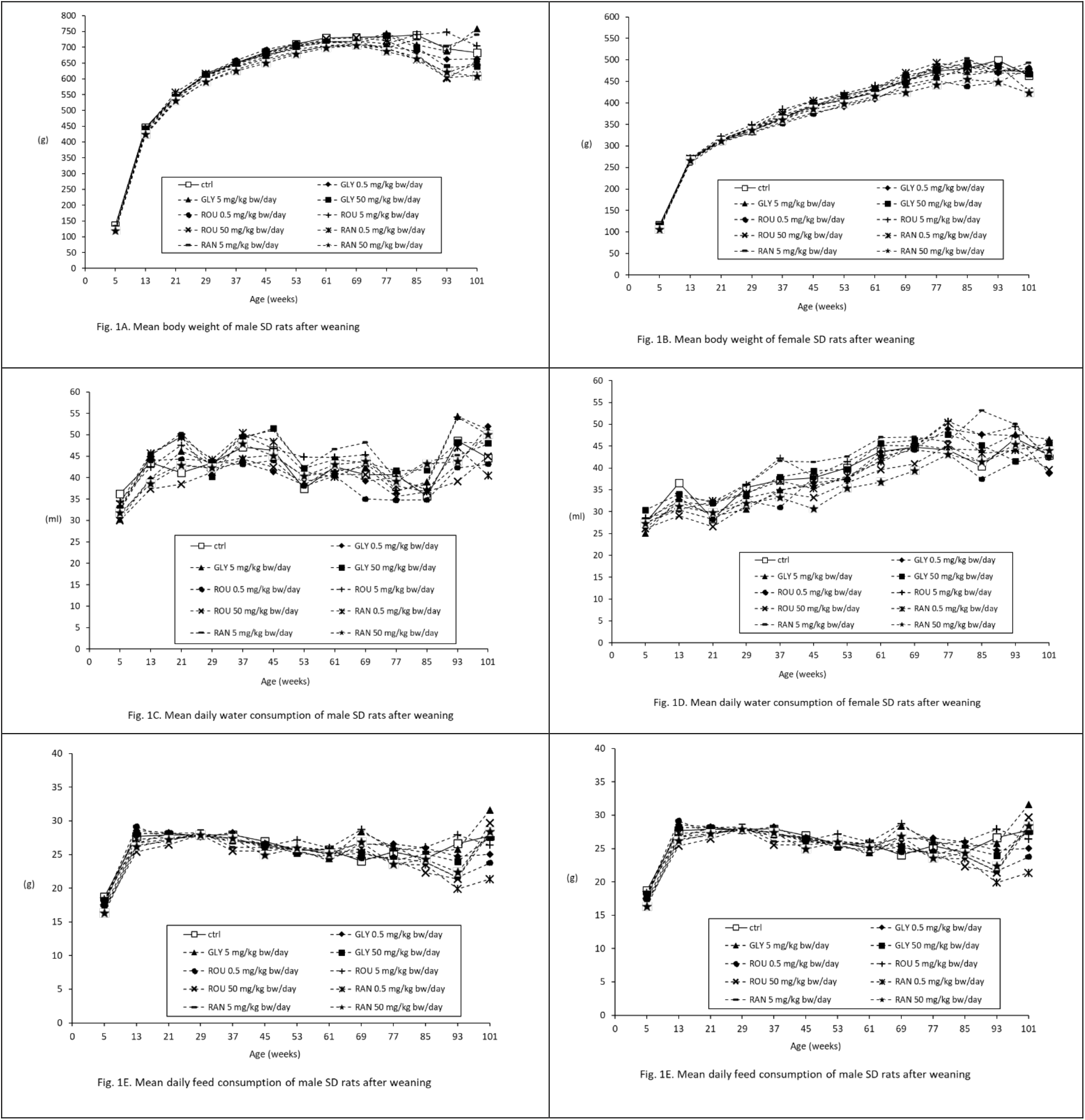
Mean body weight and mean water and feed consumption in male and female Sprague-Dawley rats after weaning.

The analysis of survival times in males and females did not show statistically significant differences between treated groups and control animals with any of the tested substances. In males treated with glyphosate, the survival functions followed a dose-related trend (p=0.0180). Graphical representations of the Kaplan-Meier curves for males and females (**Figure 2**) shows the similarity of the survival functions.

**Figure 2.**
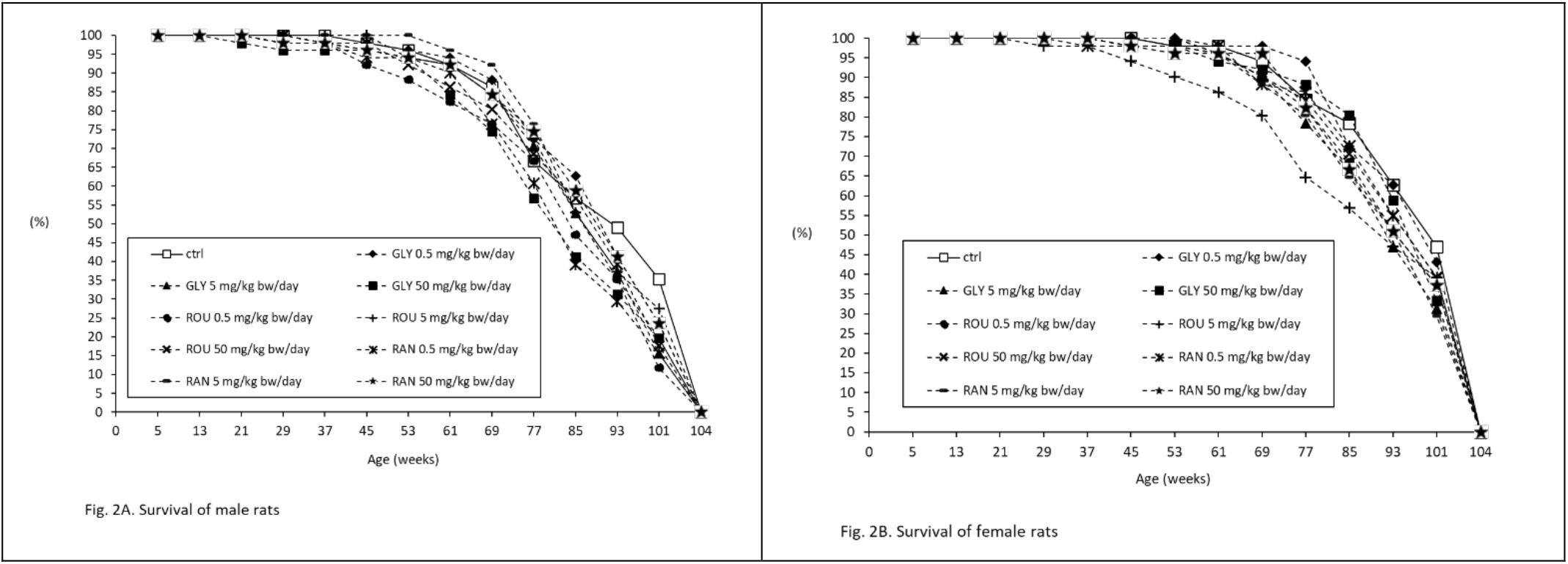
Survival in male and female Sprague-Dawley rats

In the cases of leukemia, the Tarone-Ware test showed that failure rates increase with increasing concentrations of the tested substance. Statistically significant results were observed for males, considering all substances together (p=0.0150), and in animals treated with Roundup Bioflow vs Control (p=0.0149). The poly-3 test was therefore performed to assure that the leukemia incidence analyses were correct and appropriate.

### Incidence of leukemia

Leukemia incidence following exposure to glyphosate, Roundup Bioflow or RangerPro is summarized in **Tables 1-3**.

Overall leukemia incidence in the treated groups was 1.53% (14/918 rats); 2.18% (10/459) in males and 0.87% (4/459) in females. Lymphoblastic leukemia was the most frequent subtype observed (6/14, 42.9%), followed by monocytic leukemias (5/14, 35.7%) and myeloid leukemias (3/14, 21.4%). No brothers or sisters from the same litter were affected by leukemia. No cases of leukemia were observed in untreated control animals of either sex.

#### Glyphosate

Increased incidence of lymphoblastic leukemia was observed in males exposed to pure glyphosate in a dose-dependent fashion (p=0.0419, Cochran-Armitage trend test; p=0.0335, poly-3 test). Similar results were obtained when considering males and females together (M+F, p=0.0421 Cochran-Armitage trend test; p=0.0372, poly-3 test). Two cases of monocytic leukemia were observed in male animals treated with pure glyphosate, one treated with 0.5 mg/kg bw/day and the other with 5 mg/kg bw/day. In females, one case of myeloid leukemia was observed in the group treated with the lowest dose of glyphosate (0.5 mg/kg bw/day) (**Table 1**).

#### Roundup Bioflow

A significantly increased trend in incidence of lymphoblastic leukemia was observed in both males (p=0.0419, Cochran-Armitage trend test; p=0.0425, poly-3 test) and females treated with Roundup Bioflow (p=0.0419, Cochran-Armitage trend test; p=0.0439, poly-3 test). When males and females were considered together, the statistical significance with both the Cochran-Armitage trend test (p=0.0072) and the poly-3 test (p=0.0078) became stronger.

A significantly increased trend in incidence of monocytic leukemia was observed in males treated with Roundup Bioflow (p=0.0419, Cochran-Armitage trend test; p=0.0425, poly-3 test). When males and females were considered together, statistical significance was confirmed with both tests (Cochran-Armitage trend test, p=0.0421; poly-3 test, p=0.0432).

A significantly increased trend in incidence of all types of leukemias combined was observed in males (p=0.0071, Cochran-Armitage trend test; p=0.0083, poly-3 test), and in females treated with Roundup Bioflow (p=0.0419, Cochran-Armitage trend test; p=0.0439, poly-3 test). When males and females were considered together, the statistical significance observed for all types of leukemia combined became stronger (p=0.0014, Cochran-Armitage trend test; p=0.0016, poly-3 test) (**Table 2**).

#### RangerPro

A significantly increased trend in incidence of lymphoblastic leukemia was observed in both males (p=0.0071, Cochran-Armitage trend test; p=0.0092, poly-3 test) and females treated with RangerPro (p=0.0419, Cochran-Armitage trend test; p=0.0411, poly-3 test). When males and females were considered together, the statistical significance observed with both the Cochran-Armitage trend test (p=0.0014) and the poly-3 test (p=0.0016) became stronger.

A significantly increased trend in incidence of monocytic leukemia was observed in males treated with RangerPro (p=0.0419, Cochran-Armitage trend test; p=0.0459, poly-3 test). In females treated with RangerPro, one case of monocytic leukemia was observed in an animal treated with 5 mg/kg bw/day.

Myeloid leukemia was diagnosed in one male treated with 0.5 mg/kg bw/day and in one male treated with 5 mg/kg bw/day of RangerPro.

For all types of leukemia combined, a significantly increased trend in incidence was observed in males treated with RangerPro (p=0.0313, Cochran-Armitage trend test ; p=0.0371, poly-3 test); When males and females were considered together, the statistical significance became stronger (p=0.0195, Cochran-Armitage trend test ; p=0.0218, poly-3 test) (**Table 3**).

#### Early onset of leukemia

The mean age at death among all animals bearing leukemia was 97 weeks (standard deviation ±13) for the 0.5 mg/kg bw/day groups; 68 weeks (standard deviation ±33) for the 5 mg/kg bw/day groups; and 57 weeks (standard deviation ±26) for the 50mg/kg bw/day groups (**Figure 3**).

**Figure 3.**
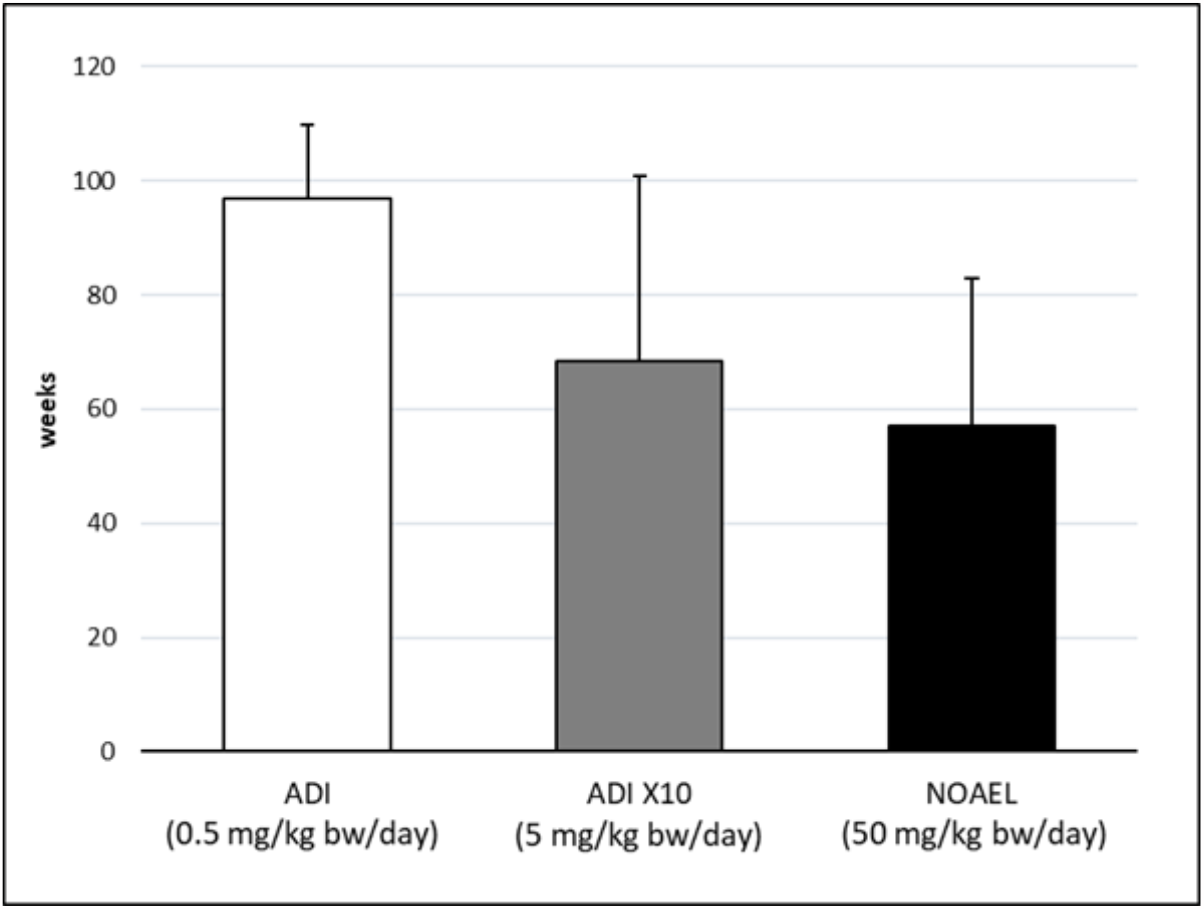
Mean (with standard deviation) weeks of age at death of animals bearing tumors, divided by doses of 0.5, 5, and 50 mg/kg body weight/day corresponding to EU Acceptable Daily Intake (ADI), ADIx10 and the EU No Observed Adverse Effect Level (NOAEL) for glyphosate. The 3 treatment groups (glyphosate, Roundup Bioflow, RangerPro) are reported combined per dose. ADI mean weeks of age 97; ADI x10 mean weeks of age 68; NOAEL mean weeks of age 57).

Notably, 43% of animals bearing leukemias (6/14) died before the first year of age (52 weeks): 5 of these 6 animals were from the highest dose groups (50 mg/kg bw/day) and 1 from the intermediate dose groups (5 mg/kg bw/day). These cases of early-onset and early death from leukemia (before 52 weeks of age) were equally distributed among the different treatment groups: 2 in the glyphosate group, 2 in Roundup Bioflow and 2 in Ranger Pro.

Leukemias are rare malignancies in SD rats. For this reason, we compared the age at death of animals bearing leukemias with the historical controls data available from the Ramazzini Institute (490 animals) (Gnudi et al. 2023) and the US National Toxicological Program (1179 animals) (NTP 2022). With a total of 1669 historical controls, only 16 animals were diagnosed with leukemia (overall incidence rate of 0.96%), with an incidence of 4/490 at the Ramazzini Institute (incidence rate of 0.82%) and an incidence of 12/1179 at the National Toxicology Program (incidence rate 1.02%). Of these16 cases, none died before the first year of age. This finding contrasts shapely with the results observed here, whereas in the present study 43% of the leukemias-related deaths in the glyphosate and GBHs treated groups occurred before the first year of age (**Figure 4**).

**Figure 4.**
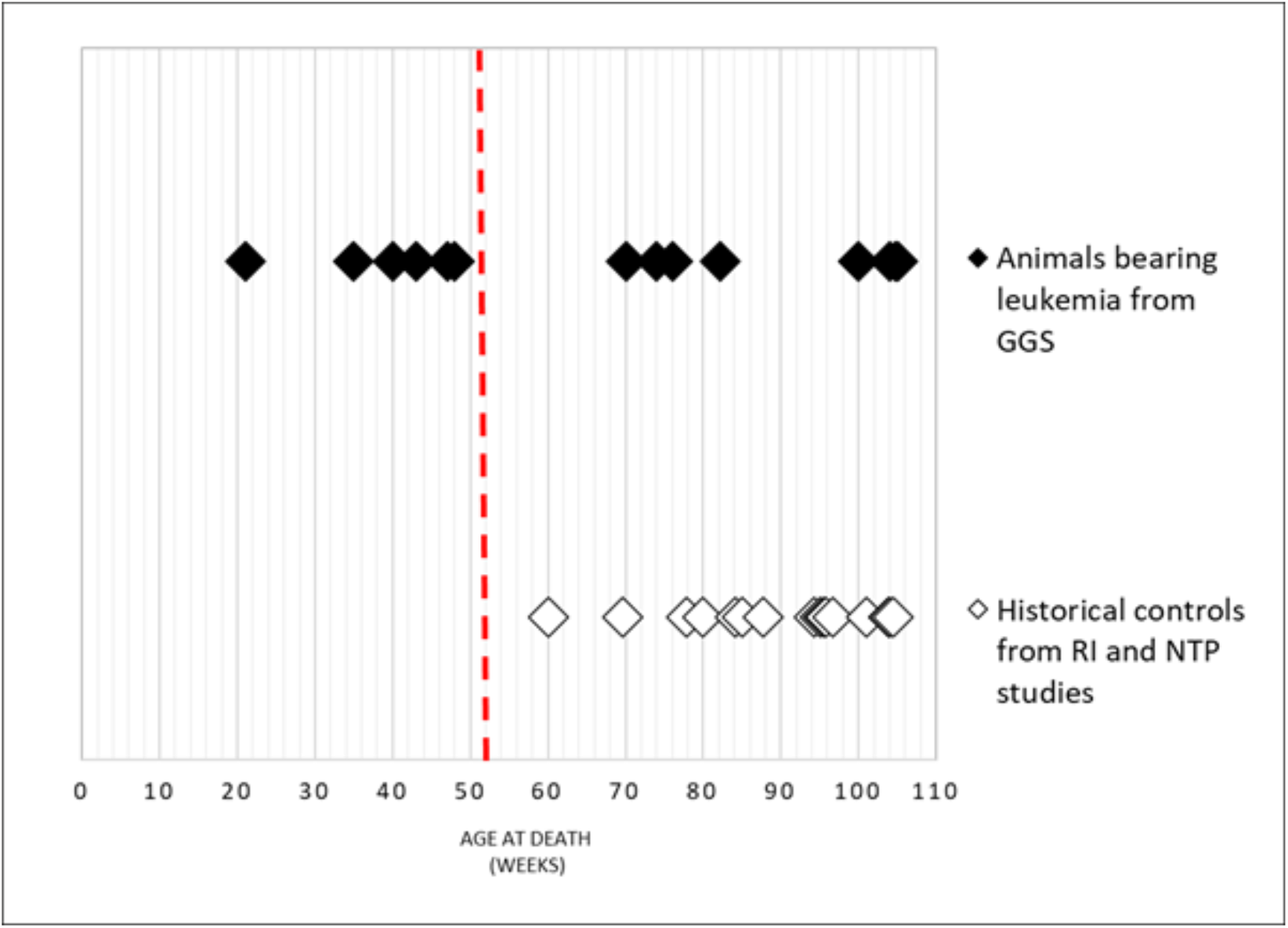
Age at death of animals (Sprague-Dawley rats) with leukemia from GGS () or from Ramazzini Institute (RI) and US National Toxicology Program (NTP) historical controls (). For GGS all animals belong to the treated groups; animals from historical controls belong only to not treated groups from various experiment. Week 52, which corresponds to 1 year of age, is dashed in red.

## Discussion

The two main findings of our study were a significant, dose-related increased trend in leukemia incidence in SD rats treated with glyphosate and GBHs at exposure levels corresponding to the EU ADI and the EU NOAEL, and early onset leukemia in exposed rats. Dose-related trends in leukemia incidence were observed in males (M) and females (F), with a majority of the significant results observed in males. When males and females were considered together (M+F), the statistical significance consistently became stronger.

In the animals exposed to glyphosate, a significantly increased trend in incidence of lymphoblastic leukemia was observed (M, M+F). In the Roundup Bioflow-treated animals, significantly increased trends in incidence of lymphoblastic leukemia (M, F, M+F), monocytic leukemia (M, M+F), total myeloid leukemia (M, M+F), and all leukemias combined (M, F, M+F) were observed. In the RangerPro-treated animals, significantly increased trends in the incidence of lymphoblastic leukemia (M, F, M+F), monocytic leukemia (M) and all leukemias combined (M, M+F) were observed. 43% of leukemia deaths in the glyphosate and GBH treated groups occurred before the first year of age (52 weeks). No case of leukemia was observed in untreated concurrent control animals.

Our results indicate that GBHs have stronger leukemogenic effects than glyphosate alone.

Interestingly, about half of the leukemia deaths observed in the rats exposed to glyphosate and GBHs in the present study occurred at less than one year of age, which is comparable to less than 35-40 years of age in humans (Sengupta 2013). By contrast, no cases of leukemia death have been observed prior to one year of age in over 1600 SD untreated control rats studied by the US National Toxicology Program (NTP) and the Ramazzini Institute over the past decade. This dose-related early onset of leukemia in the glyphosate and GBH-treated groups, is likely attributable to the fact that herbicide exposure in the present study began prenatally through treatment of pregnant dams.

Our study has limitations. One limitation is that although we compared glyphosate and its formulations (Roundup Bioflow and RangerPro), we could not explore the effects of the single adjuvants present in the GBHs. The findings on leukemia presented here are the first but not the last findings to be expected from the GGS. The evaluations of all the other neoplastic lesions that are currently ongoing will allow to elucidate the full carcinogenic potential of glyphosate and GBHs in our model.

To date, evidence from mechanistic studies in humans and animal models suggests that glyphosate and its formulations possess several of the ten key characteristics of carcinogens (Rana et al. 2023), including genotoxicity, epigenetic alteration, oxidative stress, chronic inflammation, gut microbiome perturbations, and endocrine disruption (Benbrook et al. 2023; Bukowska et al. 2022; Lesseur et al. 2021; Manservisi et al. 2019; Mao et al. 2018; Mesnage et al. 2021a; R. Mesnage et al. 2022a; R. Mesnage et al. 2022b; Truzzi et al. 2021). These studies support our finding of increased leukemia in experimental animals exposed to glyphosate and GBHs.

Our results further strengthen the “sufficient evidence of carcinogenicity in experimental animals” of the IARC evaluation and are consistent with the epidemiological evidence that has shown increases in incidence of myeloid and lymphoid malignancies in humans exposed to glyphosate and GBHs.

### Conclusions

This report from the carcinogenicity arm of the Global Glyphosate Study found that glyphosate and GBHs at exposure levels corresponding to the EU ADI and the EU NOAEL caused a statistically significant dose-related increased trend in leukemia incidence, a very rare malignancy in Sprague-Dawley rats. An additional very important finding is that about half of the leukemia deaths seen in the glyphosate and GBH groups occurred at less than one year of age, comparable to ages less than 35-40 years in humans. By contrast, no case of leukemia was observed in the first year of age in more than 1600 Sprague-Dawley historical controls in carcinogenicity studies conducted by either the Ramazzini Institute or the US National Toxicology Program.

## Acknowledgements

The authors would like to acknowledge the memory of Dr. Luciano Bua and his major contributions to this work . The authors would like to acknowledge Dr. Alberto Mantovani, former scientist at the Istituto Superiore di Sanità, for his participation and support to the Global Glyphosate Study

